# Object-Centered Spatial Learning in Dynamic Contexts: History-Driven Distractor Suppression and Target Enhancement

**DOI:** 10.1101/2024.09.04.611200

**Authors:** Yayla A. Ilksoy, Dirk van Moorselaar, Sander A. Los, Jan Theeuwes

## Abstract

The world around us is inherently structured and often repetitive. Research has shown that we can implicitly learn to prioritize relevant objects and locations while filtering out distracting information, creating an integrated priority map for attention allocation. The current study examines whether providing an object-like reference frame would induce an object-centered attentional bias or whether the bias would remain in egocentric (viewpoint-centered) coordinates. The search display consisted of six stimuli that were surrounded by a wheel and square frame. In two experiments, either a distractor or a target appeared more frequently in one location, leading to the suppression or enhancement of that location, respectively. Learning blocks were followed by test blocks, where the frame rotated, creating egocentric-matching and object-centered locations. These experiments showed that both target and distractor learning relied on an egocentric reference frame only. In follow-up experiments, the likely target and distractor location rotated dynamically with the frame during learning. This revealed that participants can learn to enhance a likely target location in an object-centered manner. We hypothesized that while space-based learning feeds into a priority map reliant on an egocentric reference frame, object-based learning allows for implicit prioritization of subparts of objects independent of their spatial orientation.

**Public significance statement:** Our brains cannot process the overwhelming amount of visual information we encounter daily. Fortunately, the world is inherently structured and has patterns we can learn, helping us to selectively focus on what’s important and ignore distractions. This study examines how we use these patterns in dynamic, ever-changing environments. The findings show that when regularities are learned in static settings, our learned attentional biases remain tied to our static viewpoints. But in dynamic environments, attentional priorities can be tied to objects, irrespective of the objects’ orientation, while suppressed distractor locations remain tied to specific viewpoints. This discovery helps us understand how our brain adapts to complex, real-world situations.

## Introduction

In our daily lives, we are constantly bombarded with a continuous stream of visual information. Yet, this does not overwhelm us, for the world is inherently structured and often repetitive. Numerous studies have demonstrated our ability to extract these regularities, known as statistical learning, which enables us to prioritize certain spatial locations with relevant information, while filtering out distracting information (Ferrante et al., 2018; Geng & Behrmann, 2002; Jiang et al., 2013; Wang & Theeuwes, 2018b; see Theeuwes et al., 2022 for a review). This type of learning has been qualified as implicit, occurring largely outside of conscious awareness and without deliberate intention to learn (Turk-Browne et al., 2005). The general concept is that selection history continuously contributes to an integrated priority map, where the weights regulate the allocation of both overt and covert attention (Awh et al., 2012; Bisley & Goldberg, 2010; Theeuwes et al., 2022). One way to study selection history effects is in paradigms in which either targets or distractors appear more often in one location of the search display than in any of the other possible locations. In case of targets this results in a spatial bias towards this location, whereas for distractors there is an attentional bias away from this location (e.g. Ferrante et al., 2018; Huang et al., 2022; Wang & Theeuwes, 2018b). This phenomenon is generally referred to as space-based learning, where it is assumed that the location of a specific object in space is attentionally suppressed or enhanced (Theeuwes et al., 2022).

If learning is space-based, it implies that the position of an object must be encoded relative to a particular frame of reference. Two primary reference frames that are typically distinguished are the egocentric and allocentric reference frames (O’Keefe, 1978). An egocentric reference frame represents the spatial coordinates of objects relative to the observer, including coordinates relative to the eyes (known as retinotopic coordinates), as well as coordinates relative to the head or body. In contrast, in an allocentric reference frame the locations of objects are encoded in relation to other objects or environmental landmarks, independent of the observer’s viewpoint. In this respect, it is noteworty that most studies on implicit statistical learning have relied on static search displays, which may not fully capture the dynamic nature of real-world environments where our perspectives constantly shift as we move around. This raises the question of whether the assumed priority map is tied to an allocentric or egocentric frame of reference.

Studies attempting to address this question, either by altering participants’ viewpoints (Jiang & Swallow, 2013, 2014) or by making eye movements (Chang & Golomb, 2024; Ilksoy et al., 2024), have mostly found evidence that statistically learned attentional biases depend on an egocentric reference frame. However, a recent study by Chang & Golomb (2024) demonstrated that spatial suppression can operate independently of an egocentric reference frame when it is learned in a dynamic context that favors spatiotopic processing (i.e., the same location within space).

Interestingly, two recent studies have shown that statistical learning does not solely modulate the priority landscape in a space-based manner but can also give rise to preferential biases within objects, independent of the objects’ orientation or location in space (van Moorselaar & Theeuwes, 2023, 2024a). Here, the spatial relationships among the parts of the object are likely not encoded relative to the observer (egocentric frame) or other external objects (allocentric “world-centered” frame). Instead, they are encoded relative to other parts of the same object, an arrangement that could be described as an “object-centered” allocentric reference frame.

Previous research on exogenous cuing, rather than statistical learning, showed that retinotopic (egocentric) and object-centered (allocentric) reference frames can operate simultaneously (Theeuwes et al., 2013). They used a design involving an abrupt onset and a rotating cross. One of the arms of a cross-shaped frame was cued with an abrupt onset. Notably, after the cross rotated, exogenous attention was directed not only towards the previously cued location on the screen (retinotopic egocentric coordinates) but also transferred to the new position of the previously cued arm (object-centered allocentric coordinates).

In the present study, we adopted a similar design to investigate whether history-driven attention exhibits comparable behavior. Specifically, we examined whether providing a consistent, object-like allocentric reference frame would induce an object-centered attentional bias or instead whether the bias would remain in egocentric coordinates. Additionally, we tested this for both target and distractor regularities. Notably, while previous studies have shown that statistical learning can occur within objects (van Moorselaar & Theeuwes, 2023, 2024a), these findings have been limited to target regularities alone. Given that prior research suggests target and distractor learning may rely on partly distinct mechanisms (Ferrante et al., 2018), and that distractor learning is often less robust (Li et al., 2023; Yu et al., 2023), it remains unclear whether distractor learning can occur within objects as target learning does.

We employed the additional singleton task used by Wang & Theeuwes (2018a) in which a distractor singleton (Experiment 1a) or target singleton (Experiment 1b) is presented more often in one location compared to all other locations. Additionally, the search display was surrounded by a frame to provide environmental landmarks and thereby allow for allocentric learning. Learning blocks were followed by test blocks, in which the frame would occasionally rotate. The question was whether the high-probability location that participants had learned to suppress (Experiment 1a) or enhance (Experiment 1b) would rotate along with the frame (allocentric location; same location with respect to environmental landmarks or “object”) or would remain in the same location as in the learning blocks (egocentric location; same location with respect to viewpoint).

## Experiment 1a

### Methods

The Ethical Review Committee of the Faculty of Behavioral and Movement Sciences of the Vrije Universiteit Amsterdam approved the present study. Sample size was predetermined based on previous studies that used similar designs (de Waard et al., 2022; Ivanov & Theeuwes, 2021; Kong et al., 2020). In these studies, the effect size (Cohen’s d) of the critical comparison between the high-probability and low-probability distractor location was > 0.7. In the current experiment we are interested in a possible transfer of the effect from one location to another. Therefore, we expect the effect size to be reduced and used a medium effect size (*d* = 0.4) in an a priori power analysis for one-tailed paired sampled t-tests. With an alpha level of .05 and a power of .80, the sufficient number of participants would be 41 (Faul et al., 2007). To counterbalance the conditions, 48 adults (22 females, M age = 28.7 years old, SD age = 6.5) were recruited for monetary compensation via the online platform Prolific (www.prolific.co; £8.79 for participation). All participants signed informed consent before the study and reported that they did not suffer from color blindness.

#### Apparatus and stimuli

We adopted a modified version of the additional singleton paradigm (Wang & Theeuwes, 2018a). The experiment was created in *OpenSesame* (Mathôt et al., 2012) and conducted in an online environment on a JATOS server (Lange et al., 2015). Because the experiment was conducted online, we describe the stimuli in terms of pixels instead of visual degrees. The search display consisted of five diamonds (subtended by 40 pixels) and one circle (radius of 30 pixels) or vice versa (see figure 1). Participants had to search for the unique shape (i.e. the target; in this example the circle). Each shape had a red (RGB: 255/0/0) or green (RGB: 0/200/2) outline and was filled on one side (left or right; randomly chosen) with the same color while the other side had the background color. The stimuli were presented on an imaginary circle with a radius of 195 pixels, centered at fixation, against a black background (RGB: 0/0/0). To provide environmental context and aid spatial orientation, the search items were presented inside a six-arm wheel (radius of 250 pixels). Each stimulus was enclosed by the ends of rectangular frames, which were interconnected at the center of the display. The stimuli and the surrounding wheel were embedded within a square outline (550 × 550 pixels). The upper side of the square was orange (RGB: 255/166/2), while the lower side of the square was striped. To further increase the availability of environmental context, the fixation cross at the center of the display was designed as a star shape consisting of three intersecting lines. One line was striped with a width of 2 pixels, whereas the other two lines were solid, with widths of 1 and 3 pixels. This visual arrangement ensured that participants could observe the rotation of the stimuli within a defined spatial context provided by the surrounding landmarks.

**Figure 1.**
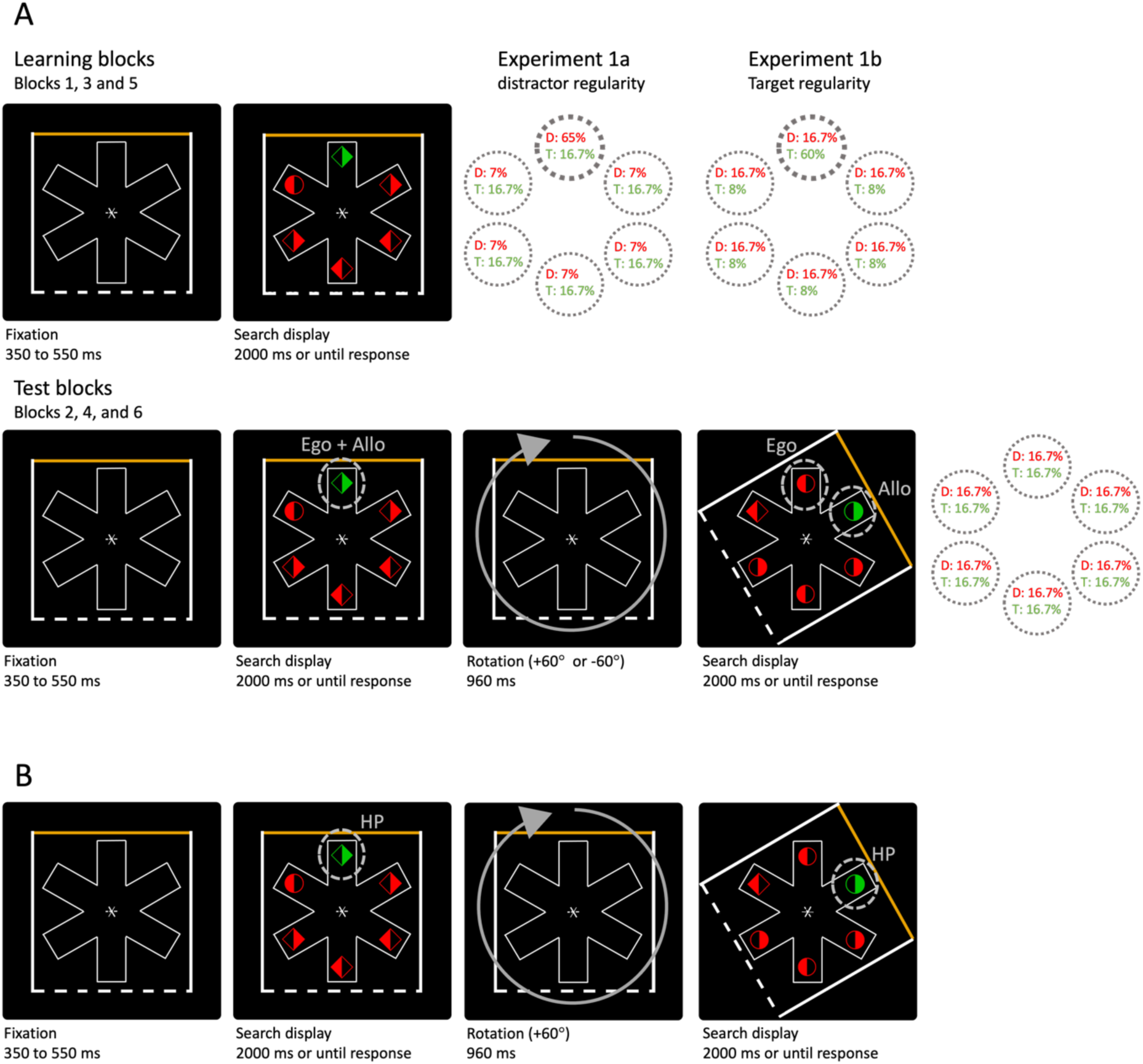
Experimental procedure. *Note.* (A) Example of a trial sequence in Experiment 1. Learning blocks (blocks 1, 3 and 5) alternated with test blocks (blocks 2, 4 and 6). Trials started with a fixation cross that was surrounded by a wheel frame embedded in a square outline (350 to 550 ms). Subsequently, a search display was presented for 2000 ms or until response. In this example, the target is the circle because it differs in shape compared to the other stimuli. The salient distractor is the green diamond. In the learning blocks, the singleton distractors (Experiment 1a) or the targets (Experiment 1b) appeared with a higher probability at one specific location (HP location; in this example the top location). In the test blocks, the singleton distractors and targets appeared at each location equally often. Every few trials (a randomly selected sequence of one to eight consecutive trials), the search display rotated either clockwise or counterclockwise (+60° or −60°). This resulted in an egocentric-matching and allocentric-matching location compared to the HP location in the learning blocks. (B) Example of a trial sequence in Experiment 2. The singleton distractors (Experiment 2a) or the targets (Experiment 2b and 2c) appeared with a higher probability at one specific location within the surrounding wheel and square frame. The search display consistently rotated clockwise (+60°) after two to eight consecutive trials and continued to rotate until the end of the block. Each time the search display rotated, the HP location rotated along with the display.

#### Procedure

Each trial started with the presentation of the fixation cross, together with the surrounding wheel and square frame, which remained visible throughout the experiment. After a randomly jittered interval (350 and 550 ms), the search display was presented for 2000 ms or until a response was made. Participants had to search for a circle among diamonds or vice versa. They were instructed to indicate via button press, as quickly and accurately as possible, whether the left (‘z’ key) or right side (‘/’ key) of the identified target was filled. If participants failed to respond or made an erroneous response, a warning message was displayed.

Search displays included a uniquely colored distractor in 66.7% of the trials. This singleton distractor shared the same shape as the other distractors but differed in color (red or green with an equal probability). The experiment consisted of different blocks. In learning blocks, singleton distractors appeared with a higher probability (65%) at one specific location (high-probability location; counterbalanced across participants) compared to the other locations (all 7%; low probability locations). In alternating order, learning blocks were followed by test blocks, in which the distractor appeared with equal probability across all display locations (16.7%). Within test blocks, every few trials (a randomly selected sequence of one to eight consecutive trials) search displays rotated either clockwise or counterclockwise. This rotation procedure allowed for a dissociation between egocentric locations (i.e. the same spatial location within the search display) and allocentric locations (i.e. the same location within the frame). Each rotation consisted of a 60° shift over a duration of 960 ms, causing each allocentric stimulus location to move either one step to the left or the right (see figure 1A). Notably, the display was never rotated by more than 60° from the initial orientation. The display was equally likely to be presented in three orientations: the initial orientation (0°), rightward tilted orientation (60°), and leftward tilted orientation (−60°). This ensured distinct positioning for egocentric and allocentric locations when the display was tilted (60° or −60°). In the 0° orientation, egocentric and allocentric locations overlapped. In both the learning and test blocks, the target was presented at each egocentric and allocentric location equally often (16.7%).

The experiment consisted of six blocks of 180 trials each. A test block always followed a learning block, resulting in three blocks of both types. Participants started with a practice block of 30 trials and repeated the practice block until they reached an accuracy of 70% or higher and were faster than 1100 ms on average. At the end of the experiment, participants were asked whether they noticed the statistical regularities and which location within the array they thought the high-probability distractor location was. Notably, these questions were interspersed with unrelated questions to avoid influencing responses to the study-related questions.

#### Statistical analysis

First, participants with an accuracy of 2.5 standard deviation from the overall accuracy were excluded and replaced. Subsequently, trials with response times (RTs) faster than 200 ms or slower than 2000 ms were excluded. Furthermore, incorrect trials were excluded for the reaction time (RT) analyses. Trials with RTs exceeding 2.5 standard deviations from the mean RT per block type per subject were excluded. Finally, participants with an RT of 2.5 standard deviation from the overall RT were also removed and replaced.

For the main analyses, only the RT results will be reported, since there was no evidence of a speed-accuracy trade-off (See Supplementary Materials for accuracy results). The RTs were subjected to repeated-measures analyses of variance (ANOVAs), followed by planned comparisons with paired-sample t-tests. Where sphericity was violated, Greenhouse-Geiser corrected p-values are reported. Effect sizes (partial eta squared or Cohen’s d) were reported for each comparison. In the case of non-significant findings, Bayes factors in support of the null hypothesis were reported (BF_01_). First, the RTs in the learning blocks were analyzed to confirm that the participants had successfully learned to suppress the high-probability (HP) distractor location. This effect was characterized by faster RTs when distractors appeared at the HP location compared to the average of all other locations (the low-probability location). To verify that the observed effect was not attributed to inter-trial priming effects (Maljkovic & Nakayama, 1996) but rather to long-term statistical learning, the analysis was repeated exclusively for trials where the distractor did not appear at the same location as in the preceding trial (distractor priming trials). Second, the RTs in the test blocks were analyzed to determine whether this suppression effect remained at the egocentric location (i.e., at the same spatial location on the screen) following the rotation of the search display or if it was transferred to the allocentric location (i.e., the same location within the surrounding wheel and square frame). The latter would imply that the suppression dynamically rotated along with the rotation of the wheel and the surrounding square frame. In the test blocks, the low-probability (LP) location included all stimulus positions when the frame was oriented at −60° or 60°, except for the egocentric and allocentric locations. Additionally, we collapsed the trials immediately following rotation and repeated our main analysis on these trials alone to determine whether the observed effects emerged immediately after rotation.

Two factors of awareness of the HP location were considered. First, participants were categorized as subjectively aware if they answered that they noticed that the distractor was more frequently in one location compared to all others. Second, objective awareness of the HP location was examined by calculating the distance between the selected HP location and the actual HP location for each subject (including the subjectively unaware participants). A t-test was conducted to determine whether the average distance significantly differed from chance level. If the t-test indicated that, on average, participants correctly identified the HP location, the main analyses were repeated, incorporating an interaction between the Distractor condition and participants’ awareness (correct or incorrect selection of the HP location) as a between-subjects factor to determine whether awareness influenced the observed effect. In case of non-significant findings, Bayes factors will be provided that represent the comparison between repeated-measures ANOVAs that contain the interaction and those that do not. The BF_excl_ indicates how much more likely the data are under a model without the interaction compared to one with the interaction.

#### Transparancy and openness

All data and analysis codes are available at https://osf.io/8r2ut/. Data were analyzed using R, version 4.3.1 (R Core Team, 2018) and Bayes factors were determined using JASP version 0.17.1 (Lange et al., 2015) All data was collected in 2024. This study’s design and its analysis were not pre-registered. The participant pool contains a balanced number of male and female participants. Additionally, collecting data online allowed us to include participants from diverse global backgrounds. This, together with the fact that the study’s focus is on fundamental cognitive processes, suggests limited constraints on generality.

## Results

In total, 11 participants were excluded and replaced based on their accuracy (eight participants) and their RTs (three participants). In the final sample, exclusion of incorrect responses (6.8%) and data trimming (2.7%) resulted in an overall loss of 9.5% of the trials.

### Learning blocks

A repeated-measures ANOVA with Distractor condition (HP location, LP location and no distractor) as a within-subject factor revealed a significant main effect on mean RTs (F (2, 94) = 212.22, *p* < .001, 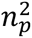 = .82; see Figure 2A). Subsequent planned comparisons showed that relative to no distractor trials, RTs were slower when the distractor appeared at the HP or LP location (all *t*’s > 13.4 and *p*’s < .001). Crucially, RTs were faster when the distractor appeared at the HP location compared to the LP location (*t* (47) = 9.41, *p* < .001, *d* = 1.36), indicative of learned attentional suppression at the HP distractor location. Notably, these results remained consistent after excluding distractor priming trials (*t* (47) = 9.53, *p* < .001, *d* = 1.38), indicating that the observed suppression effect was not solely driven by intertrial priming.

**Figure 2.**
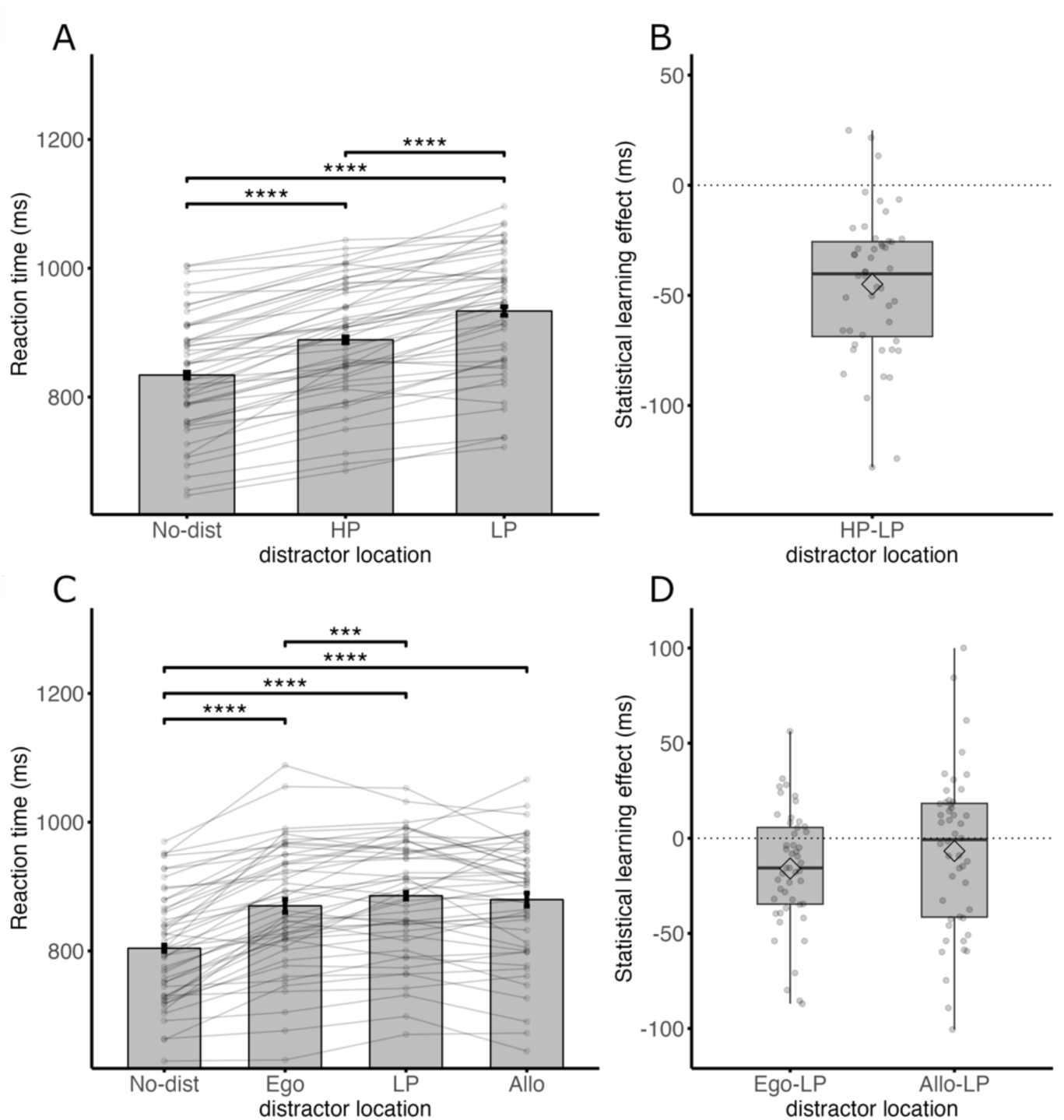
Learned attentional suppression in Experiment 1a. *Note.* (A and B) Mean response times in the learning blocks. Participants responded faster when the distractor was presented at the HP compared to the LP location. (C and D) Mean response times in the test blocks. Participants responded faster when the distractor was presented at the egocentric location compared to the LP location. No difference was found between the allocentric location and the LP location. The height of each bar (A and C) reflects the sample average and each dot that is interconnected by a line reflects data from each individual participant. Error bars represent 95% within-subject confidence intervals (Morey, 2008). The significance bars represent the planned comparisons with paired-sample t-tests. The diamonds inside the boxplots (B and D) represent the mean difference scores and the horizontal lines represent the median difference scores.

Of the 48 participants, 14 reported noticing that the distractor appeared more frequently in one location than in others (subjectively aware). Among these, only one participant correctly identified the HP distractor location. When all participants (both subjectively aware and unaware) were asked to identify the HP distractor location, they selected a location that was, on average, 1.4 units away from the actual HP location, a deviation that did not significantly deviate from chance level (chance level: 1.5; *t* (47) = 0.59, *p* = .56, *d* = 0.085, BF_01_ = 5.4).

### Test blocks

After having established that participants learned to suppress the HP distractor location, we focused on examining the effects of visual display rotation on this learned suppression: would the suppressed location persist at the egocentric location or remap to the allocentric location? A repeated-measures ANOVA with Distractor condition (egocentric location, allocentric location, LP location and no distractor) as a within-subject factor revealed a significant main effect on mean RTs (F (2.5, 117.56) = 79.011, *p* < .001, 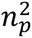 = 0.63; see Figure 2C). Subsequent planned comparisons confirmed that, compared to no distractor, mean RTs were slower when the distractor was presented at the egocentric, allocentric or LP location (all *t*’s > 10.8 and *p*’s < .001). Importantly, compared to the LP location, mean RTs were faster when the distractor was presented at the egocentric location (*t* (47) = 3.55, *p* < .001, *d* = 0.51), whereas there were no significant differences between the LP location and the allocentric location (*t* (47) = 1.11, *p* = .27, *d* = 0.16, BF_01_ = 3.6). This pattern of results was already present in the first trial following the rotation of the frame (Ego vs. LP: *t* (47) = 2.79, *p* < .01, *d* = 0.40; Allo vs. LP: *t* (47) = 1.40, *p* = .17, *d* = 0.20, BF_01_ = 3.6).

## Discussion

The current findings are in line with previous studies that showed that history-driven suppression is learned in egocentric coordinates (Ilksoy et al., 2024; Jiang & Swallow, 2013, 2014). The addition of a stable object-like allocentric reference frame did not induce an attentional bias that was independent of viewpoint and environmentally stable. In Experiment 1b, we investigated whether the same holds when participants learn to prioritize a likely target location.

## Experiment 1b

Experiment 1b was identical to Experiment 1a but now with a HP target location instead of a HP distractor location. Forty-eight adults (21 females, M age = 27.5 years old, SD age = 8.8) were recruited for monetary compensation via the online platform Prolific (www.prolific.co; £8.25). In the learning blocks, the target appeared with a higher probability of 60% at one specific location (counterbalanced across participants). All other locations had a chance of 8% to contain a target. The distractor appeared at each location equally often (16.7%). For the main analyses, the RTs were subjected to a repeated-measures ANOVA. The within-subject factors were Target Condition (HP and LP locations in the learning blocks; egocentric, allocentric, and LP locations in the test blocks) and Distractor Presence (present, absent). Subsequently, planned comparisons were performed with paired-sample t-tests. Similar to Experiment 1, the main analysis was repeated excluding those trials where the target was presented at the same location as the previous trial (target priming trials).

## Results

In total, 15 participants were excluded and replaced based on their accuracy (nine participants) and based on their RTs (six participants). In the final sample, exclusion of incorrect responses (6.8%) and data trimming (3.2%) resulted in an overall loss of 10% of the trials.

### Learning blocks

A repeated-measures ANOVA on mean RTs revealed a significant main effect for Target condition (HP and LP location; F (1, 47) = 104.28, *p* < .001, 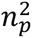 = .69; see Figure 3A), Distractor presence (present and absent; F (1, 47) = 401.57, *p* < .001, 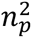 = .90), and a significant interaction between these two factors (F (1, 47) = 4.75, *p* = .034, 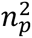 = .092). Subsequent planned comparisons showed that RTs were faster when the target was presented at the HP location compared to the LP location, both in the distractor present (*t* (47) = 8.82, *p* < .001, *d* = 1.27) and in the distractor absent condition (*t* (47) = 9.82, *p* < .001, *d* = 1.42). RTs remained faster at the HP compared to the LP location after exclusion of target priming trials (*t* (47) = 11.02, *p* < .001, *d* = 1.59).

**Figure 3.**
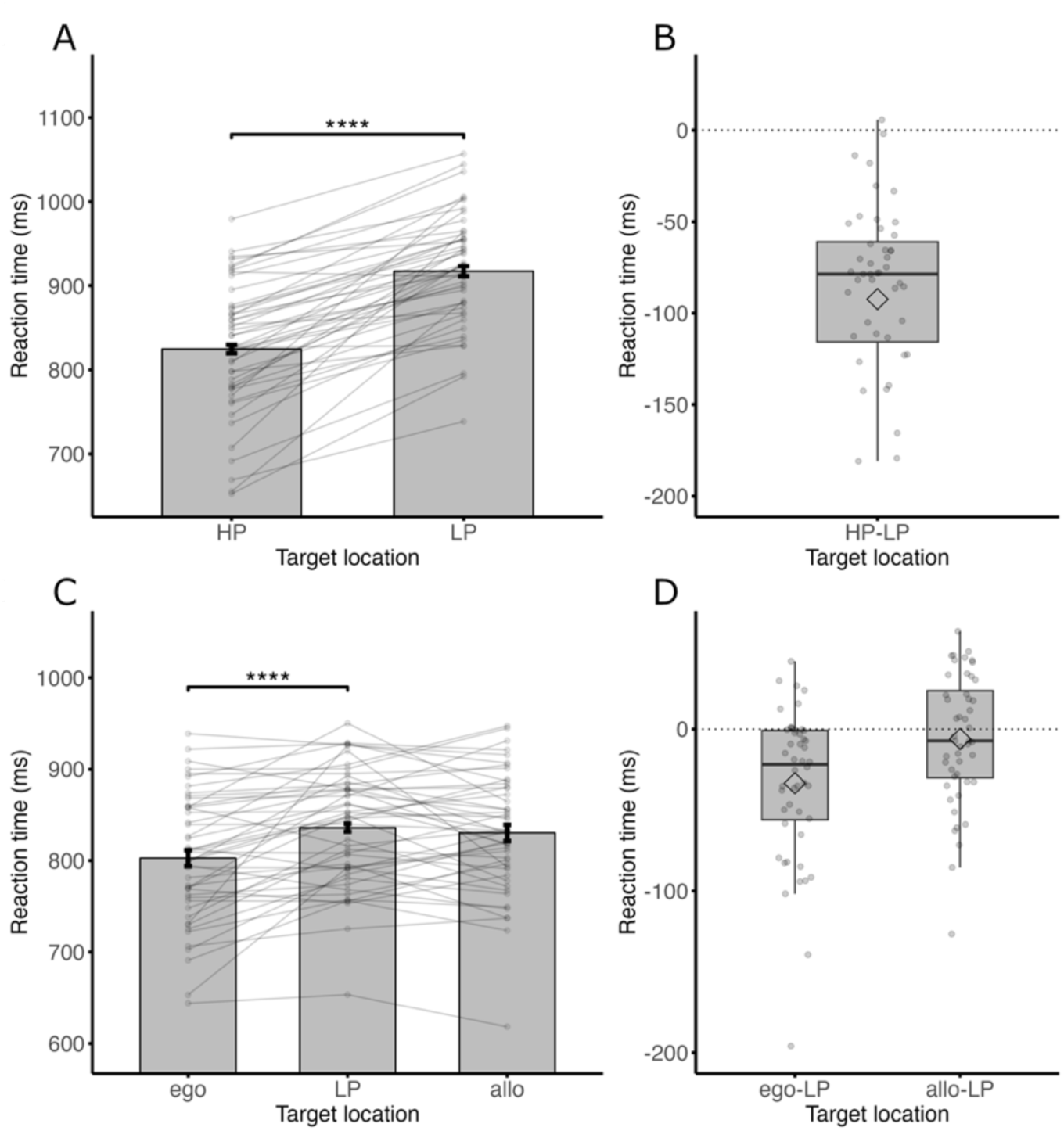
Learned attentional enhancement in Experiment 1b. *Note.* (A and B) Mean response times in the learning blocks. Participants responded faster when the distractor was presented at the HP compared to the LP location. (C and D) Mean response times in the test blocks. Participants responded faster when the distractor was presented at the egocentric location compared to the LP location. No difference was found between the allocentric location and the LP location.

Of the 48 participants, 40 reported noticing that the target appeared more frequently in one location than in others (subjectively aware). Among these, 31 participants correctly identified the HP distractor location. When all participants (both subjectively aware and unaware) were asked to identify the HP distractor location, their selected location was, on average, 0.5 units away from the actual HP location, a deviation that significantly differed from chance level (chance level: 1.5; *t* (47) = 7.51, *p* < .001, *d* = 1.10). A follow-up repeated-measures ANOVA showed that an interaction between objective awareness (i.e. correct/incorrect selection of HP target location of all participants) and Target condition was trending towards significance (RT: F (1, 46) = 3.60, *p* = .064, 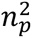 = 0.73, BF_excl_ = 3.0). However, both groups (objectively aware: 35 participants; objectively unaware: 13 participants) displayed a clear prioritizing of the HP location compared to the LP location (objectively aware: *t* (34) = 9.76, *p* < .001, *d* = 1.65; objectively unaware: *t* (12) = 7.86, *p* < .001, *d* = 2.18).

### Test blocks

A Repeated-measures ANOVA on mean RTs revealed a significant main effect for Target condition (egocentric location, allocentric location and LP location; F (2, 94) = 16.56, p < .001, 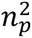= 0.26; see figure 4C) and for Distractor Presence (present, absent; F (1, 47) = 287.50, p < .001, 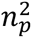= 0.86). An interaction between Target condition and Distractor presence was not significant (F (2, 94) = 0.16, *p* = .85, 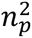 = .003, BF_excl_ = 12.0). Further planned comparisons revealed that compared to the LP location, RTs were faster when the target was presented at the egocentric location compared to the LP location (*t* (47) = 5.01, *p* < .001, *d* = 0.72), while there was no difference in RTs between the allocentric and LP location (*t* (47) = 1.044, *p* = .30, *d* = 0.15, BF_01_ = 3.8). This pattern of results manifested as early as in the first trial following the rotation of the frame (Ego vs. LP: *t* (47) = 4.30, *p* < .001, *d* = 0.62; Allo vs. LP: *t* (47) = 0.62, *p* = .54, *d* = 0.090, BF_01_ = 5.3). A follow-up analysis revealed that an interaction between objective awareness (i.e. correct/incorrect selection of HP target location of all participants) and Target condition did not reach significance (F (2, 92) = 0.90, *p* = .41, 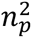 = .019, BF_excl_ = 3.3).

**Figure 4.**
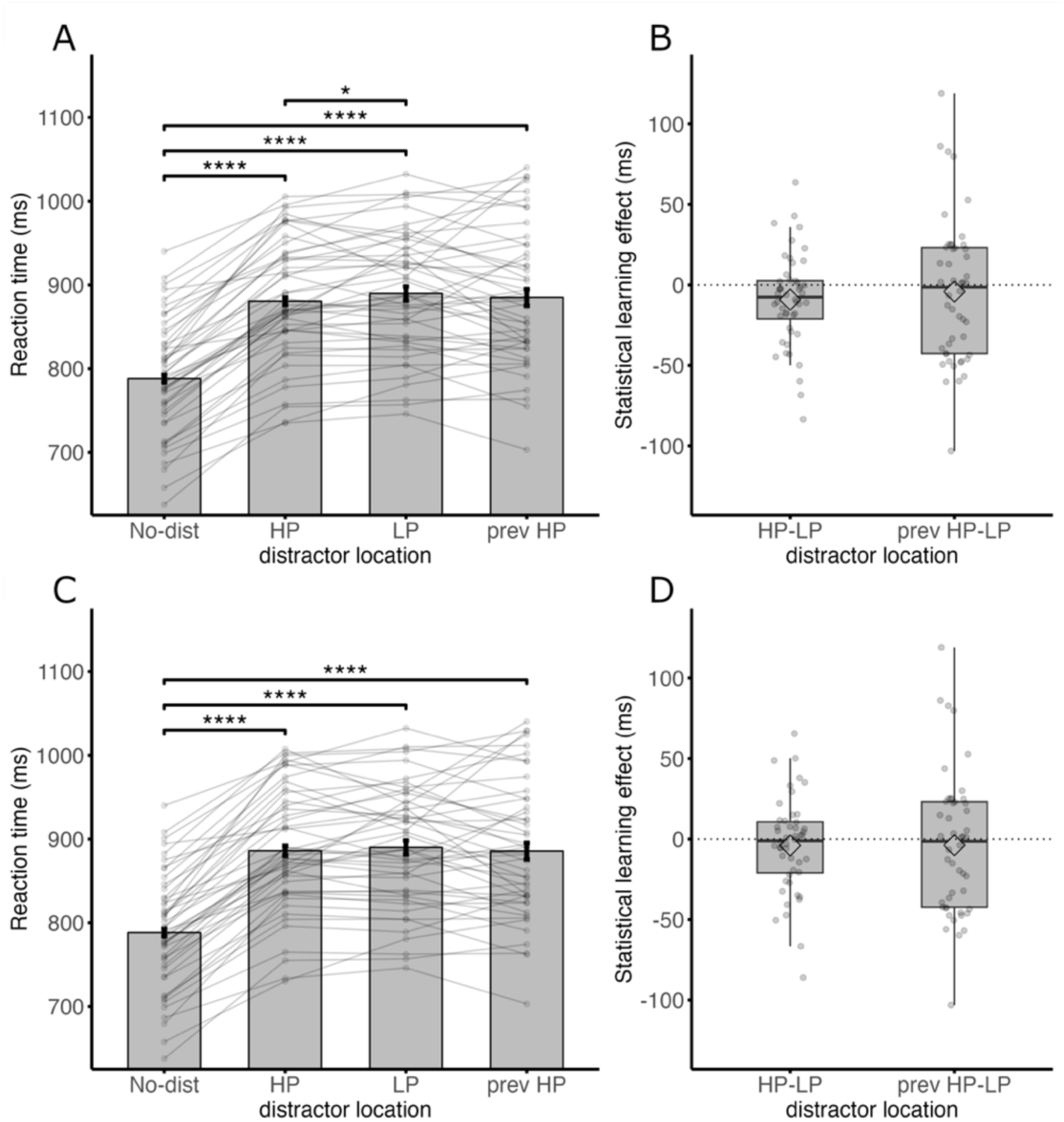
Learned attentional suppression in Experiment 2a. *Note.* (A and B) Mean response times including all trials. Participants responded faster when the distractor was presented at the HP compared to the LP location. (C and D) Mean response times excluding distractor priming trials. The difference between the HP and LP condition disappeared after excluding distractor priming trials.

## Discussion

Experiment 1b showed that, like distractors, enhancement of the probable target location was dependent on an egocentric reference frame and not on an allocentric reference frame. In other words, when the wheel and square frame rotated, the enhanced location did not rotate along with the frame but stayed in the same location in space. However, it should be noted that the spatial imbalance was only present in the static learning blocks, and it is thus possible that an allocentric attentional bias is only formed in a dynamic rather than a static environment, similar to what was suggested by Chang & Golomb (2024) regarding spatiotopic suppression.

To test this hypothesis, we performed an additional experiment in which the search display rotated from the start of the experiment. Importantly, the likely distractor location (Experiment 2a) and the likely target location (Experiment 2b and 2c) rotated along with the outer frame. The question was whether participants were able to suppress or enhance the HP location when this location is no longer static, but defined relative to the surrounding object.

## Experiment 2a

The goal of Experiment 2a was to determine whether participants could learn to suppress the HP distractor location when the HP location dynamically rotated along with the surrounding wheel and square frame. Forty-eight adults (25 females, M age = 27.4 years old, SD age = 6.1) were recruited for monetary compensation via the online platform Prolific (www.prolific.co; £5.75 for participation). Experiment 2a was identical to Experiment 1a except for the following changes.

Experiment 2a did not include test blocks (as in Experiment 1a), but only learning blocks in which the uniquely colored distractor had a high probability of ∼57% of being presented at one specific allocentric location (i.e., the same location within the wheel and square frame) ^2^. All other allocentric locations had a probability of ∼7% to contain the uniquely colored distractor. Importantly, across an experimental block, the uniquely colored distractor appeared equally often at each egocentric location (i.e., the spatial locations on the screen).

In Experiment 2a, the search display consistently rotated clockwise and continued to rotate until the end of the block. The display rotated after a randomly selected sequence of trials (two to eight consecutive trials). As in the learning blocks of the previous experiment, the HP condition contains more distractor priming trials, in which the distractor appears at the same location as in the previous trial, compared to the LP condition. However, in contrast to the previous experiment, there is not one physical HP location, but the HP location rotates along with the outer frame. Consequently, the locations that were previously the HP location might be affected by a lingering inter-trial priming effect. Therefore, the LP condition was divided into two distinct conditions: the three locations that were in the clockwise direction (LP condition) and the two locations that were in the counterclockwise direction (previous HP condition).

The experiment consisted of five blocks of 144 trials, preceded by a practice block of 15 trials. Participants responded to the side of the target that was filled by pressing the left arrow button for left and the right arrow button for right. Due to technical issues, the answers to the question about awareness of the HP distractor location were not registered.

## Results

In total, 14 participants were excluded and replaced based on their accuracy (eight participants) and their RTs (six participants). In the final sample, exclusion of incorrect responses (6%) and data trimming (3.7%) resulted in an overall loss of 9.7% of the trials.

A repeated-measures ANOVA with Distractor condition (no distractor, HP location, LP location or previous HP location) as within-subject factor revealed a significant main effect on mean RTs (F (2.12, 99.63) = 151.06, *p* < .001, 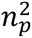 = .76; see Figure 4). Subsequent planned comparisons showed that relative to no distractor trials, RTs were slower when the distractor appeared at the HP, LP or previous HP location (all *t*’s > 12.9 and *p*’s < .001). Critically, RTs were faster when the distractor was presented at the HP location compared to the LP location (*t* (47) = 2.23, *p* = .030, *d* = 0.32). However, the difference disappeared after excluding distractor priming trials (*t* (47) = 0.94, *p* = .35, *d* = 0.14, BF_01_ = 4.2; see figure 4C and 4D). Moreover, no significant differences were found between the previous HP location and the HP location (*t* (47) = 1.04, *p* = .30, *d* = 0.15, BF_01_ = 3.8) or LP location (*t* (47) = 0.65, *p* = .52, *d* = 0.094, BF_01_ = 5.2).

## Discussion

The findings show that participants were able to learn to suppress the HP distractor location when the location dynamically rotated along with its environment. However, it is crucial to recognize that the effect appears to be relatively small (∼9 ms), especially when compared to previous findings that used the same paradigm with static displays. Even considering the lower probability in Experiment 2 compared to Experiment 1, one would not expect such a significant reduction in the suppression effect (Lin et al., 2021). Furthermore, the difference was no longer present after excluding distractor priming trials. Therefore, it is uncertain whether the effect can be related to long-term statistical learning.

## Experiment 2b

The goal of Experiment 2b was to see whether participants could learn to enhance the HP target location when the HP location dynamically rotated along with the surrounding wheel and square frame. Forty-eight adults (24 females, M age = 26.5 years old, SD age = 5.8) were recruited for monetary compensation via the online platform Prolific (www.prolific.co; £5.63 for participation). Experiment 2b was identical to Experiment 2a except for the following changes. The target had a higher probability of ∼50% to be presented at one specific allocentric location. All other allocentric locations had a probability of ∼10% to contain the target. Importantly, across an experimental block, the target was presented at each egocentric location equally often. The experiment consisted of 4 blocks of 162 trials each.

## Results

In total, nine participants were excluded and replaced based on their accuracy (five participants) and based on their RTs (four participants). In the final sample, exclusion of incorrect responses (5.8%) and data trimming (3.4%) resulted in an overall loss of 9.2% of the trials.

A repeated-measures ANOVA with Target condition (HP location, LP location and previous HP location) and Distractor condition (present, absent) as within-subject factors revealed a main effect for Target condition (F (2, 94) = 6.52, *p* < .01, 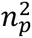 = 0.12), as well as a main effect for Distractor condition (F (1,47) = 422.80, *p* < .001, 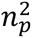 = 0.90). The interaction between Target condition and Distractor condition was not significant (F (1.71, 80.4) = 0.65, *p* = .52, 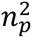 = 0.014, BF_excl_ = 3.0). Paired-sample t-tests showed faster RTs when the target was presented at the HP location compared to the LP location (*t* (47) = 7.50, *p* < .001, *d* = 1.082; see figure 5A) and compared to the previous HP location (*t* (47) = 5.057, *p* < .001, *d* = 0.73). These differences in RTs were still present after excluding target priming trials (HP-LP: *t* (47) = 6.56, *p* < .001, *d* = 0.95; HP-prev HP: *t* (47) = 3.90, *p* < .001, *d* = 0.56). No significant differences were found between the previous HP location and the LP location (*t* (47) = 1.21, *p* = 0.23, *d* = 0.17, BF_01_ = 3.2). After repeating the analysis by including only the trials immediately following rotation and excluding target priming trials, we found no significant main effect of the Target condition (F (2, 94) = 1.019, *p* = .37, 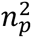 = 0.021, BF_01_ = 10.1).

**Figure 5.**
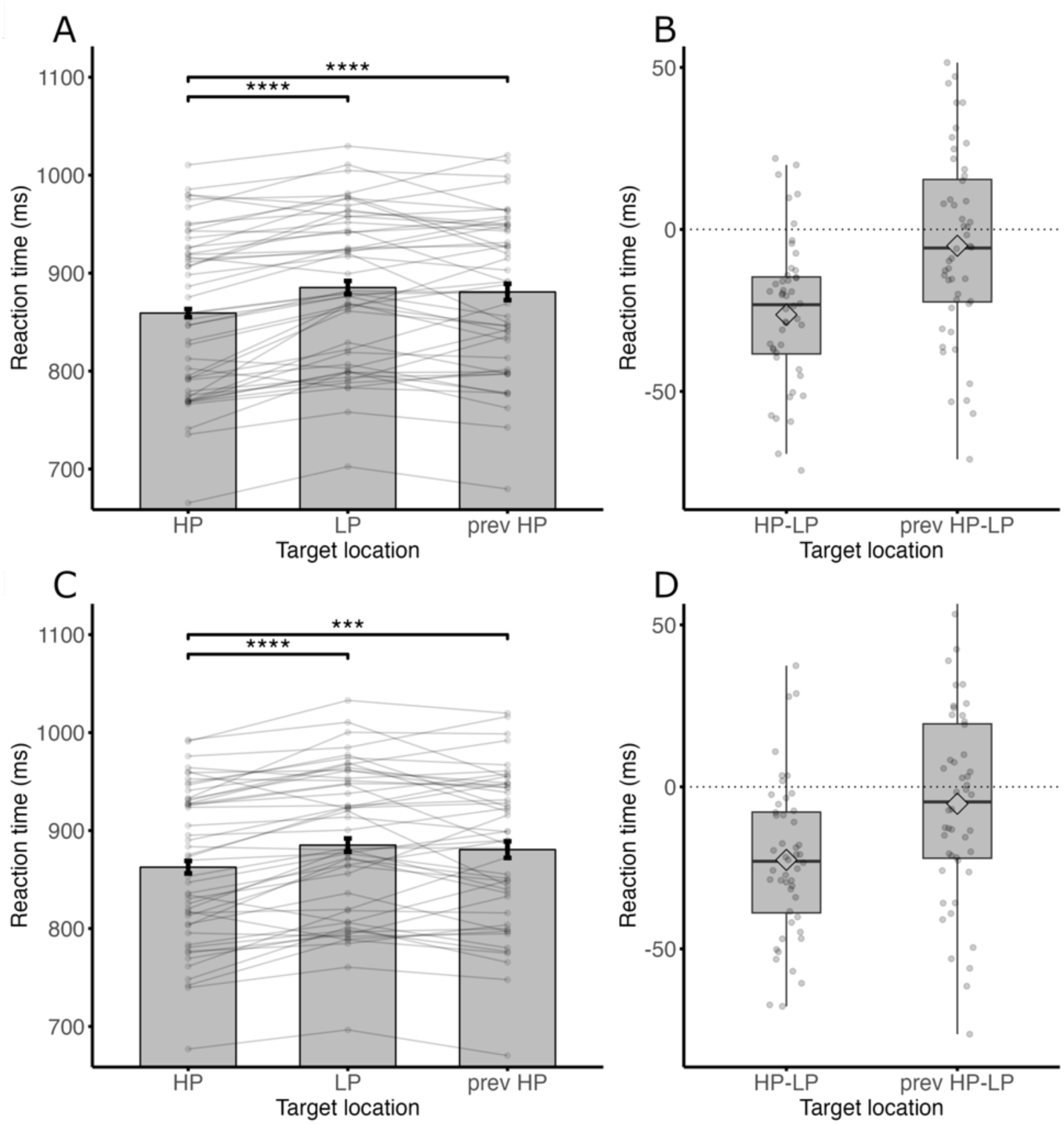
Learned attentional enhancement in Experiment 2b. *Note.* (A and B) Mean response times including all trials. Participants responded faster when the target was presented at the HP compared to the LP location and compared to the previous HP location. (C and D) Mean response times excluding target priming trials. The differences between the HP and LP condition and HP and previous HP condition were still significant after excluding target priming trials.

Although the results appear to be in line with allocentric learning of target regularities, after careful consideration of the data, it became apparent that the decreased RTs in the HP condition relative to the LP condition might be explained by the fact that HP target conditions contained fewer distractor present displays compared to the LP condition.^3^ This imbalance likely accounts for a significant portion of the reduced RTs observed in the HP condition compared to the LP condition.

## Discussion

The main findings of experiment 2b seem to suggest that an attentional bias towards a likely target location may depend on an object-centered allocentric reference if the attentional bias is formed in a dynamic setting. This conclusion is premature however, given that the effect may have been influenced by the higher proportion of distractor displays in the LP location compared to the HP location, rather than by the statistical regularities alone. Therefore, we repeated the experiment without any colored distractors. If the RTs are again faster in the HP condition compared to the LP condition, it cannot be attributed to distractor presence and solely to statistical learning.

## Experiment 2c

Experiment 2c was identical to Experiment 2b with the only difference that there were no colored distractors presented in the whole experiment. Forty-eight adults (24 females, M age = 30.0 years old, SD age = 7.4) were recruited for monetary compensation via the online platform Prolific (www.prolific.co; £5.25 for participation). Awareness of the HP target location was assessed at the end of the experiment by asking participants whether they noticed the statistical regularities regarding the target and whether they could indicate which location within the surrounding frame was the HP target location. Awareness of the HP target location was examined by calculating the distance between the selected HP location and the actual HP location for each subject. Additionally, because Experiment 2b revealed that the learning effect was not significant immediately following rotation, we divided the trials into two separate conditions – the first three trials following rotation (first half) and the subsequent three trials (second half) – and included this factor in our main analysis.

## Results

In total, 11 participants were excluded and replaced based on their accuracy (four participants) and based on their RTs (seven participants). In the final sample, exclusion of incorrect responses (4%) and data trimming (3.3%) resulted in an overall loss of 7.3% of the trials.

After excluding target priming trials (See Supplementary table 3 for results including target priming trials), a repeated measures ANOVA with Target condition (HP location, LP location or previous HP location) and Post-rotation condition (first half, second half) as within-subject factors revealed a main effect for Target condition (F (2, 94) = 16.96, *p* < .001, 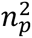 = .27), as well as a main effect for Post-rotation condition (F (1, 47) = 49.94, *p* < .001, 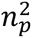 = .52; see figure 6). Notably, the interaction between Target condition and Post-rotation condition was also significant (F (2, 94) = 22.71, *p* < .001, 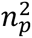 = .33). Paired-sample t-tests indicated that in the first three trials after rotation (first half), RTs were faster in the previous HP condition compared to both the HP (*t* (47) = 2.038, *p* = .047, *d* = 0.29) and LP (*t* (47) = 3.62, *p* < .001, *d* = 0.52) conditions, whereas in the subsequent three trials (second half), RTs were faster in the HP condition compared to the previous HP condition (*t* (47) = 5.97, *p* < .001, *d* = 0.86) and the LP condition (*t* (47) = 6.92, *p* < .001, *d* = 1.00)^4^. There was no significant difference between the HP and LP conditions in the first three trials after rotation (*t* (47) = 1.03, *p* = .31, *d* = 0.15, BF_01_ = 3.9), nor between the previous HP and LP conditions in the subsequent three trials after rotation (*t* (47) = 0.23, *p* = .82, *d* = 0.089, BF_01_ = 6.2).

**Figure 6.**
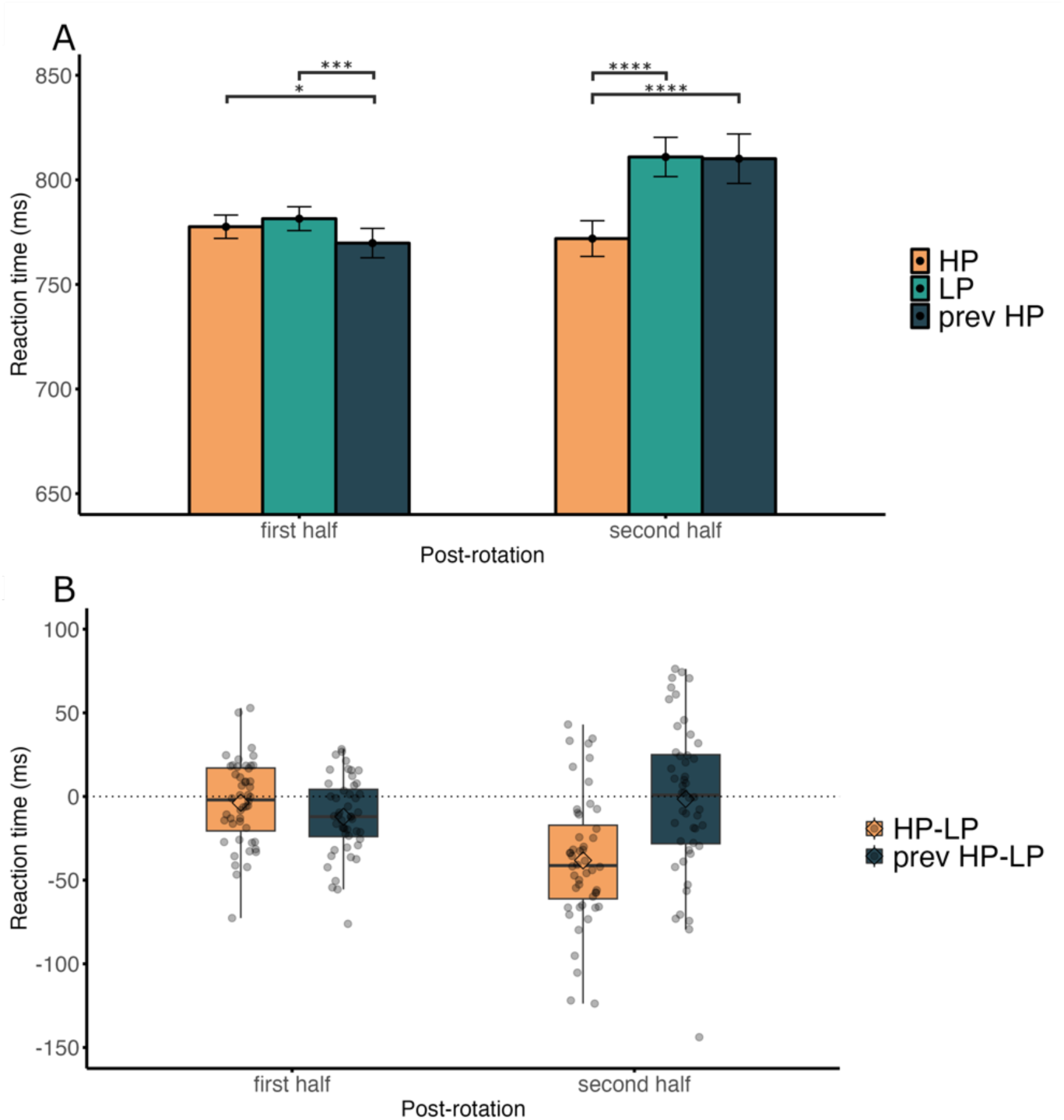
Learned attentional enhancement in Experiment 2c. *Note.* (A and B) Mean response times excluding target priming trials. In the first three trials following the rotation (first half), participants responded faster when the target appeared in the previous HP location compared to both the current HP and LP locations. However, in the subsequent three trials, participants responded faster when the target appeared in the current HP location compared to the previous HP and LP locations.

Of the 48 participants, 31 reported noticing that the target appeared more frequently in one location within the frame than in others (subjectively aware). Among these, 5 participants correctly identified the HP target location. Furthermore, analysis of the awareness scores indicated that participants (both subjectively aware and unaware) chose a location that was on average 1.5 units away from the actual HP target location, which did not significantly differ from chance level (chance level: 1.5; *t* (47) = 0.32, *p* = .75, *d* = 0.046, BF_01_ = 6.1).

## Discussion

Again, it was shown that participants can learn to enhance a likely target location that dynamically rotates along with its allocentric reference frame. Additionally, the findings suggest that the attentional bias was learned implicitly. This effect cannot be attributed to the increased presence of a salient distractor in the HP location, as was the concern in Experiment 2b.

Interestingly, the data indicate that following the rotation of the allocentric frame, the attentional bias was gradually updated from the previous HP location to the current HP location. One plausible explanation for the gradual shift in attentional bias after rotation is cumulative location priming effects over time. However, we controlled for immediate priming effects (i.e., n-1, n-2), making this explanation less likely. Another plausible explanation is that the learning system requires time to reset and reconfigure the priority settings, reflecting a genuine long-term learning effect.

## General discussion

Recent work has shown that attention is not only shaped by top-down and bottom-up processes but also by statistical regularities in the visual field (Ferrante et al., 2018; Geng & Behrmann, 2002; Jiang et al., 2013; Wang & Theeuwes, 2018b; see Theeuwes et al., 2022 for a review). Locations with a high probability of distractors are attentionally suppressed while locations with a high probability of targets are attentionally enhanced. Notably, most studies that have focused on visual statistical learning have used static displays, in which the search display did not move in space. The real world, however, is dynamic and we constantly observe the world from different perspectives. In the current study, we aimed to test whether providing a consistent, object-like allocentric reference frame would induce an object-centered attentional bias or, as shown with static displays, whether the bias would remain in egocentric coordinates. Experiment 1 showed that when a spatial imbalance is learned in a static search display, the spatial bias remains in egocentric coordinates once the display starts to rotate. This finding was observed for both distractor and target learning. In contrast, Experiment 2 revealed that an object-centered bias can emerge when the imbalance is learned in a dynamic context; however, this effect was specific to target learning.

The absence of object-centered attention in experiment 1 could be explained by space-based learning overshadowing object-based learning, primarily because the stationary display encountered at the start of the experiment did not necessitate an object centered approach for efficient target selection. Consequently, once space-based learning was adopted, participants continued to depend on this learning system during the test phase when the search display started to rotate. Within this perspective, it is not surprising that we observed only egocentric suppression and egocentric enhancement in Experiment 1, as previous research suggests that space-based learning relies on an egocentric reference frame. For example, Jiang & Swallow (2013, 2014) conducted a series of experiments which revealed that learned attentional biases towards a likely target location moved along with the participants’ viewpoint when participants walked around the search display. Additionally, when the spatial bias remained in the same location, but participants moved around the display, they were unable to learn to prioritize that location.

Our findings are also consistent with a study by Chang & Golomb (2024), where participants learned to suppress a likely distractor location in a learning array and then made a single switch in fixation to an unbiased test array. In this context, the learned suppression was maintained in retinotopic coordinates rather than remapped to (environmentally stable) spatiotopic coordinates (see also Ilksoy et al., 2024). However, when participants dynamically shifted their fixation between arrays, while maintaining the spatial imbalance, suppression occurred in spatiotopic instead of retinotopic coordinates. While the previous study differs from the current one in that it does not focus on an object-centered reference frame and specifically addresses distractor learning, both studies highlight that implicit biases are typically based on an egocentric reference frame unless formed in a dynamic context, in which case they can become stable in environmental coordinates.

So, while space-based statistical learning seems to feed into a priority map that relies on an egocentric reference frame, prioritized parts of an object can be coded relative to the rest of the object, independent of its orientation or location in space (van Moorselaar & Theeuwes, 2023, 2024a). It makes sense that biases within objects are tied to coordinates that are independent of viewpoint, since objects constantly move around in space. The spatial location of an object, however, is crucial for actions like grabbing it, which explains why these locations are coded relative to the viewer.

The inability to adjust a location bias in response to a viewpoint change corresponds to the finding that space-based statistical learning is independent of context (Britton & Anderson, 2019; de Waard et al., 2022) and generalizes across tasks (Jiang, Swallow, Won, et al., 2014; van Moorselaar & Theeuwes, 2024b). This implies that implicit statistical learning depends on an inflexible system that, once a bias is established, remains unaffected by changes in environmental or task conditions (but also see, de Waard et al., 2023; Gao et al., 2023). This rigidity is particularly striking given that implicit biases tend to persist even after spatial imbalances are corrected (Ferrante et al., 2018; Jiang et al., 2013). While participants can acquire new biases when exposed to new regularities, pre-existing biases only diminish gradually (Jiang et al., 2013; Wang & Theeuwes, 2020). Taken together, this implies that space-based statistical learning results in an inflexible priority map where previously learned priorities continue to exist in egocentric coordinates and may co-occur with newly learned biases.

The asymmetry between space-based statistical learning and object-centered statistical learning suggests that they rely on different neural mechanisms. Space-based statistical learning could be resolved in early visual areas, which are known to be retinotopically organized, and can be achieved by altering synaptic weights. Indeed, decreased brain activity in response to distractor regularities is already evident at the earliest stages of visual processing (Richter et al., 2024; Zhang et al., 2022). Object-based statistical learning, on the other hand, must rely, at least in part, on higher brain areas, since it cannot rely solely on egocentric retinotopic coordinates. Indeed, already in 1993, Goodale & Milner (1992) proposed two distinct neural pathways: the dorsal stream for processing spatial information and the ventral stream for object recognition.

Further evidence supporting these distinct mechanisms comes from the findings of Experiment 2c. The object-centered bias toward the likely target location only gradually remapped after the object’s rotation, whereas the egocentric bias in Experiment 1 was immediately evident in the first trial following rotation. This supports the idea that spatial biases are proactively modulated (Duncan et al., 2023; Ferrante et al., 2023; Huang et al., 2021, 2022, 2023), enabling suppression or enhancement of specific locations in space in anticipation of upcoming stimuli. However, when priorities are tied to an object frame, they must be updated reactively after the object’s presentation. Thus, a plausible explanation for the findings in Experiment 2c is that the learning system required time to reset after the object’s frame rotated, reactively reconfiguring the priority settings.

The current study not only highlights a discrepancy between space-based learning and object-based learning but also reveals a distinction between implicit distractor learning and implicit target learning. In Experiment 2, the effects of distractor learning were smaller compared to target learning and disappeared entirely once distractor priming trials were excluded. One explanation is that distractor learning takes longer to establish, and that extended exposure to the spatial imbalance might eventually result in distractor learning as well. However, Experiment 2a (distractors) already included one additional block compared to Experiment 2b and 2c (targets). Moreover, no significant interaction between Block number and Distractor condition was found (see Supplementary Figures 1-5 for RTs across blocks), suggesting that the bias remained stable across blocks. Therefore, we do not consider it likely that additional trials would have resulted in distractor learning. An alternative explanation is that while target regularities within objects can be dynamically learned, distractor regularities cannot.

Ferrante et al. (2018) already proposed that target and distractor learning might partly rely on different mechanisms. It is commonly observed that suppressing a likely distractor location reduces the efficiency of target selection in that location, while enhancing a likely target location increases interference from distractors. While such indirect effects might initially suggest that target and distractor learning operate through similar mechanisms, Ferrante et al. (2018) revealed that the indirect effect of target learning on distractor suppression is stronger than the reverse. This asymmetry supports the idea that target and distractor learning rely, at least partially, on distinct mechanisms. Furthermore, while participants are able to respond faster to target locations that are predicted by the location of the target on the previous trial (Li & Theeuwes, 2020; Yu et al., 2023), participants do not display any corresponding sequential learning with predictable distractor locations (Li et al., 2023; Yu et al., 2023). Together with the current study, this suggests that distractor learning has a less widespread influence compared to target learning.

Another important factor to consider when discussing the reference frames of history-driven learning is the role of explicit knowledge of the spatial imbalances in the search display. One of the few studies to demonstrate allocentric-dependent statistical learning also found that this learning was associated with greater explicit awareness of the likely target location (Jiang, Won, et al., 2014). While explicit knowledge of spatial imbalances does not appear to influence implicitly formed egocentric biases, it can still guide attention toward allocentric locations (Jiang, Swallow, & Sun, 2014). However, in the current study, the difference in findings between Experiment 1b (egocentric bias) and Experiment 2c (object-centered bias) is unlikely to be attributable to differences in explicit knowledge. In both experiments, most participants were unable to identify the correct high-probability target location. If anything, participants in Experiment 1b, where explicit knowledge would be expected to play a smaller role, appeared to have slightly more awareness of the spatial imbalance than those in Experiment 2c. This suggests that explicit knowledge is unlikely to be related to the observed biases in our findings.

Unfortunately, we did not record awareness data for Experiment 2a, leaving us unable to determine whether and how explicit knowledge of the likely distractor location influenced attentional processing. Based on the findings of Experiment 1, however, we expect that participants were less aware of the likely distractor location than the likely target location. This possible difference in the level of awareness might explain the differing results between distractor learning and target learning in Experiment 2. While this remains a plausible explanation, we do not believe awareness played a major role in inducing object-centered learning, since participants in experiment 2c demonstrated low levels of awareness of the likely target location and still enhanced the likely target location in an object-centered manner. Moreover, prior studies on statistical learning have not found an association between explicit knowledge and learning either (Gao & Theeuwes, 2022), nor was such an association found in studies on regularities within objects (van Moorselaar & Theeuwes, 2023, 2024a).

In conclusion, the current study demonstrates that the reference frame of statistical learning is shaped by the context in which spatial regularities are learned. When learning occurs in static displays, space-based learning dominates, resulting in egocentric biases that persist even when the display becomes dynamic. In contrast, object-centered learning can emerge when regularities are learned in dynamic contexts, particularly for target selection.

## Acknowledgments

This research was supported by a NWO Open competition grant 406.21.GO.034 and by a European Research Council (ERC) advanced grant 833029 – [LEARNATTEND].^1^

## Footnotes

1 The authors declare no competing financial interests.

2 Due to a programming mistake, the HP location had a lower probability in Experiment 2 compared to Experiment 1. Based on prior research, a probability of 57% is more than sufficient for a significant suppression effect (Lin et al., 2021). Across most trials, the HP location contained the distractor ∼65% of the time. However, in the final trial before rotation, the HP location contained the distractor only about 7% of the time, resulting in an overall probability of 57%. Average RTs increased with the number of trials following rotation with RTs being highest in the last trial before the next rotation. Consequently, the RTs in the LP trials were artificially increased due to a greater proportion of LP trials occurring in the final trial before rotation. To eliminate this bias, the last trial before each rotation was excluded from analysis.

3 To ensure that any observed effects in the HP target location were attributable to increased target probability rather than decreased distractor probability, distractors were presented equally across all locations. However, to maintain this equal distribution, distractors were more often presented at the HP target location when the target appeared at an LP target location. As a result, the LP condition included a higher proportion of distractor-present trials than the HP condition.

4 Given that inter-trial priming effects can persist across multiple trials (Maljkovic & Nakayama, 1996), we conducted an additional control analysis. This analysis excluded not only trials where the target appeared in the same location as the previous trial (n-1 priming) but also trials where the target appeared in the same location as two trials prior (n-2 priming). Even with these exclusions, the analysis still showed a significant difference between the HP and LP conditions (t (47) = 4.87, *p* < .001, *d* = 0.70) and the HP and previous HP conditions (t (47) = 3.76, *p* < .001, *d* = 0.54) in the second half after rotation.

## References

Awh, E., Belopolsky, A. V., & Theeuwes, J. (2012). Top-down versus bottom-up attentional control: a failed theoretical dichotomy. Trends in Cognitive Sciences, 16(8), 437–443. 10.1016/J.TICS.2012.06.010

Bisley, J. W., & Goldberg, M. E. (2010). Attention, intention, and priority in the parietal lobe. Annual Review of Neuroscience, 33, 1–21. 10.1146/ANNUREV-NEURO-060909-152823

Britton, M. K., & Anderson, B. A. (2019). Specificity and Persistence of Statistical Learning in Distractor Suppression. 10.1037/xhp0000718

Chang, S., & Golomb, J. (2024). From the eye to the world: Spatial suppression is primarily coded in retinotopic coordinates but can be learned in spatiotopic coordinates. 10.31234/OSF.IO/THMP7

de Waard, J., Bogaerts, L., van Moorselaar, D., & Theeuwes, J. (2022). Surprisingly inflexible: Statistically learned suppression of distractors generalizes across contexts. Attention, Perception, and Psychophysics, 84(2), 459–473. 10.3758/S13414-021-02387-X/FIGURES/8

de Waard, J., van Moorselaar, D., Bogaerts, L., & Theeuwes, J. (2023). Statistical learning of distractor locations is dependent on task context. Scientific Reports 2023 13:1, 13(1), 1–12. 10.1038/s41598-023-38261-z

Duncan, D. H., van Moorselaar, D., & Theeuwes, J. (2023). Pinging the brain to reveal the hidden attentional priority map using encephalography. Nature Communications 2023 14:1, 14(1), 1–13. 10.1038/s41467-023-40405-8

Faul, F., Erdfelder, E., Lang, A. G., & Buchner, A. (2007). G*Power 3: A flexible statistical power analysis program for the social, behavioral, and biomedical sciences. Behavior Research Methods, 39(2), 175–191. 10.3758/BF03193146/METRICS

Ferrante, O., Patacca, A., Di Caro, V., Della Libera, C., Santandrea, E., & Chelazzi, L. (2018). Altering spatial priority maps via statistical learning of target selection and distractor filtering. Cortex, 102, 67–95. 10.1016/J.CORTEX.2017.09.027

Ferrante, O., Zhigalov, A., Hickey, C., & Jensen, O. (2023). Statistical Learning of Distractor Suppression Downregulates Prestimulus Neural Excitability in Early Visual Cortex. Journal of Neuroscience, 43(12), 2190–2198. 10.1523/JNEUROSCI.1703-22.2022

Gao, Y., de Waard, J., & Theeuwes, J. (2023). Learning to suppress a location is configuration-dependent. Attention, Perception, and Psychophysics, 1–8. 10.3758/S13414-023-02732-2/FIGURES/2

Gao, Y., & Theeuwes, J. (2022). Learning to suppress a location does not depend on knowing which location. Attention, Perception, and Psychophysics, 84(4), 1087–1097. 10.3758/S13414-021-02404-Z/FIGURES/4

Geng, J. J., & Behrmann, M. (2002). Probability Cuing of Target Location Facilitates Visual Search Implicitly in Normal Participants and Patients with Hemispatial Neglect. 10.1111/1467-9280.00491 13(6), 520–525. https://doi.org/10.1111/1467-9280.00491

Goodale, M. A., & Milner, A. D. (1992). Separate visual pathways for perception and action. Trends in Neurosciences, 15(1), 20–25. 10.1016/0166-2236(92)90344-8

Huang, C., Donk, M., & Theeuwes, J. (2022). Proactive Enhancement and Suppression Elicited by Statistical Regularities in Visual Search. Journal of Experimental Psychology: Human Perception and Performance, 48(5), 443–457. 10.1037/XHP0001002

Huang, C., Donk, M., & Theeuwes, J. (2023). Attentional suppression is in place before display onset. Attention, Perception, and Psychophysics, 85(4), 1012–1020. 10.3758/S13414-023-02704-6/FIGURES/2

Huang, C., Vilotijević, A., Theeuwes, J., & Donk, M. (2021). Proactive distractor suppression elicited by statistical regularities in visual search. Psychonomic Bulletin and Review, 28(3), 918–927. 10.3758/S13423-021-01891-3/FIGURES/3

Ilksoy, Y. A., Moorselaar, D. van, Wang, B., Los, S. A., & Theeuwes, J. (2024). Distractor suppression operates exclusively in retinotopic coordinates. BioRxiv, 2024.02.01.578407. 10.1101/2024.02.01.578407

Ivanov, Y., & Theeuwes, J. (2021). Distractor suppression leads to reduced flanker interference. Attention, Perception, and Psychophysics, 83(2), 624–636. 10.3758/S13414-020-02159-Z/FIGURES/8

Jiang, Y. V., & Swallow, K. M. (2013). Spatial reference frame of incidentally learned attention. Cognition, 126(3), 378–390. 10.1016/J.COGNITION.2012.10.011

Jiang, Y. V., & Swallow, K. M. (2014). Changing viewer perspectives reveals constraints to implicit visual statistical learning. Journal of Vision, 14(12). 10.1167/14.12.3

Jiang, Y. V., Swallow, K. M., Rosenbaum, G. M., & Herzig, C. (2013). Rapid acquisition but slow extinction of an attentional bias in space. Journal of Experimental Psychology. Human Perception and Performance, 39(1), 87. 10.1037/A0027611

Jiang, Y. V., Swallow, K. M., & Sun, L. (2014). Egocentric coding of space for incidentally learned attention: Effects of scene context and task instructions. Journal of Experimental Psychology: Learning Memory and Cognition, 40(1), 233–250. 10.1037/A0033870

Jiang, Y. V., Swallow, K. M., Won, B. Y., Cistera, J. D., & Rosenbaum, G. M. (2014). Task specificity of attention training: the case of probability cuing. Attention, Perception, and Psychophysics, 77(1), 50–66. 10.3758/S13414-014-0747-7/FIGURES/13

Jiang, Y. V., Won, B. Y., Swallow, K. M., & Mussack, D. M. (2014). Spatial reference frame of attention in a large outdoor environment. Journal of Experimental Psychology. Human Perception and Performance, 40(4), 1346–1357. 10.1037/A0036779

Kong, S., Li, X., Wang, B., & Theeuwes, J. (2020). Proactively location-based suppression elicited by statistical learning. PLOS ONE, 15(6), e0233544. 10.1371/JOURNAL.PONE.0233544

Lange, K., Kühn, S., & Filevich, E. (2015). "Just Another Tool for Online Studies” (JATOS): An Easy Solution for Setup and Management of Web Servers Supporting Online Studies. PLOS ONE, 10(6), e0130834. 10.1371/JOURNAL.PONE.0130834

Li, A. S., Bogaerts, L., & Theeuwes, J. (2023). No evidence for spatial suppression due to across-trial distractor learning in visual search. Attention, Perception, and Psychophysics, 85(4), 1088–1105. 10.3758/S13414-023-02667-8/FIGURES/5

Li, A. S., & Theeuwes, J. (2020). Statistical regularities across trials bias attentional selection. Journal of Experimental Psychology: Human Perception and Performance, 46(8), 860– 870. 10.1037/XHP0000753

Lin, R., Li, X., Wang, B., & Theeuwes, J. (2021). Spatial suppression due to statistical learning tracks the estimated spatial probability. Attention, Perception, and Psychophysics, 83(1), 283–291. 10.3758/S13414-020-02156-2/FIGURES/6

Maljkovic, V., & Nakayama, K. (1996). Priming of pop-out: II. The role of position. Perception and Psychophysics, 58(7), 977–991. 10.3758/BF03206826/METRICS

Mathôt, S., Schreij, D., & Theeuwes, J. (2012). OpenSesame: An open-source, graphical experiment builder for the social sciences. Behavior Research Methods, 44(2), 314–324. 10.3758/S13428-011-0168-7/FIGURES/4

Morey, R. D. (2008). Confidence Intervals from Normalized Data: A correction to Cousineau (2005). Tutorials in Quantitative Methods for Psychology, 4(2), 61–64. 10.20982/TQMP.04.2.P061

O’Keefe, J. (1978). The hippocampus as a cognitive map. https://www.cmor-faculty.rice.edu/~cox/neuro/HCMComplete.pdf

R Core Team. (2018). A language and environment for statistical computing. R Foundation for Statistical Computing, Vienna, Austria. Available Online: Www.R-Project.Org/(Accessed on 11 September 2020). https://scholar.google.com/scholar?hl=nl&as_sdt=0%2C5&q=R+Core+Team.+%282020%29.+A+language+and+environment+of+statistical+computing.+R+Foundation+for+Statistical+Computing.&btnG=

Richter, D., Moorselaar, D. van, & Theeuwes, J. (2024). Proactive distractor suppression in early visual cortex. BioRxiv, 2024.04.03.587747. 10.1101/2024.04.03.587747

Theeuwes, J., Bogaerts, L., & van Moorselaar, D. (2022). What to expect where and when: how statistical learning drives visual selection. Trends in Cognitive Sciences, 26(10), 860–872. 10.1016/J.TICS.2022.06.001

Theeuwes, J., Mathôt, S., & Grainger, J. (2013). Exogenous object-centered attention. Attention, Perception, and Psychophysics, 75(5), 812–818. 10.3758/S13414-013-0459-4/FIGURES/4

Turk-Browne, N. B., Jungé, J. A., & Scholl, B. J. (2005). The automaticity of visual statistical learning. Journal of Experimental Psychology: General, 134(4), 552–564. 10.1037/0096-3445.134.4.552

van Moorselaar, D., & Theeuwes, J. (2023). Statistical Learning Within Objects. Psychological Science, 34(4), 501–511. 10.1177/09567976231154804/FORMAT/EPUB

van Moorselaar, D., & Theeuwes, J. (2024a). Spatial transfer of object-based statistical learning. Attention, Perception, and Psychophysics, 86(3), 768–775. 10.3758/S13414-024-02852-3/FIGURES/2

van Moorselaar, D., & Theeuwes, J. (2024b). Transfer of statistical learning between tasks. Journal of Experimental Psychology: Human Perception and Performance. 10.1037/XHP0001216

Wang, B., & Theeuwes, J. (2018a). How to inhibit a distractor location? Statistical learning versus active, top-down suppression. Attention, Perception, and Psychophysics, 80(4), 860–870. 10.3758/S13414-018-1493-Z/FIGURES/5

Wang, B., & Theeuwes, J. (2018b). Statistical Regularities Modulate Attentional Capture. Journal of Experimental Psychology: Human Perception and Performance, 44(1), 13–17. 10.1037/XHP0000472

Wang, B., & Theeuwes, J. (2020). Implicit attentional biases in a changing environment. Acta Psychologica, 206, 103064. 10.1016/J.ACTPSY.2020.103064

Yu, H., Allenmark, F., Müller, H. J., & Shi, Z. (2023). Asymmetric Learning of Dynamic Spatial Regularities in Visual Search: Robust Facilitation of Predictable Target Locations, Fragile Suppression of Distractor Locations. Journal of Experimental Psychology: Human Perception and Performance, 49(5), 709–724. 10.1037/XHP0001120

Zhang, B., Weidner, R., Allenmark, F., Bertleff, S., Fink, G. R., Shi, Z., & Müller, H. J. (2022). Statistical Learning of Frequent Distractor Locations in Visual Search Involves Regional Signal Suppression in Early Visual Cortex. Cerebral Cortex, 32(13), 2729–2744. 10.1093/CERCOR/BHAB377

