## Supplementary Materials for "Object-Centered Spatial Learning in Dynamic Contexts: History-Driven Distractor Suppression and Target Enhancement"

### Supplementary table 1

*Descriptive results for the error rates across experiments 1 and 2*

| Experiment | Block type | Condition | Mean error rate[a] | SD |
|---|---|---|---|---|
| 1a | | | | |
| | Learning | HP | 7.45 | 31.89 |
| | Learning | LP | 10.29 | 36.89 |
| | Learning | No-dist | 5.23 | 27.03 |
| | Test | Ego | 5.01 | 24.43 |
| | Test | Allo | 6.56 | 27.48 |
| | Test | LP | 7.16 | 28.50 |
| | Test | No-dist | 3.96 | 21.74 |
| 1b | | | | |
| | Learning | HP | 5.52 | 32.00 |
| | Learning | LP | 9.19 | 40.22 |
| | Test | Ego | 5.72 | 26.52 |
| | Test | Allo | 7.01 | 29.03 |
| | Test | LP | 6.29 | 27.71 |
| 2a | | | | |
| | . | HP | 6.94 | 29.00 |
| | . | Prev HP | 6.92 | 29.08 |
| | . | LP | 7.35 | 29.78 |
| | . | No-dist | 3.13 | 20.09 |
| 2b | | | | |
| | . | HP | 5.54 | 27.87 |
| | . | Prev HP | 5.74 | 28.30 |
| | . | LP | 6.07 | 28.91 |
| 2c | | | | |
| | . | HP | 3.41 | 22.17 |
| | . | Prev HP | 3.78 | 23.26 |
| | . | LP | 4.48 | 25.14 |

[a] Proportion of errors (%)

**Supplementary table 2**

*Repeated-measures ANOVA results for the error rates across experiments 1 and 2*

| Experiment | Block type | df | F | *p*-value | *p* < .05 | Partial $\eta^2$ |
|---|---|---|---|---|---|---|
| 1a | | | | | | |
| | Learning | 2, 94 | 57.34 | < .001 | *** | 0.55 |
| | Test | 2.4, 112.7 | 9.59 | < .001 | *** | 0.17 |
| 1b | | | | | | |
| | Test | 2, 94 | 3.30 | .041 | * | 0.066 |
| 2a | | | | | | |
| | . | 2.61, 122.55 | 21.33 | < .001 | *** | 0.31 |
| 2b | | | | | | |
| | . | 1.58, 74.4 | 0.039 | .93 | n.s. | 0.00083 |
| 2c | | | | | | |
| | . | 1.6, 75.1 | 4.46 | .021 | * | 0.087 |

*Note.* Significant effects are marked as follows: *p < .05, **p < .01, ***p < .001, and non-significant results are indicated by "n.s."

**Supplementary table 3**

*Planned paired-sample t-test results for the error rates across experiments 1 and 2*

| Experiment | Block type | Planned comparison | Mean VSL effect[a] | df | *t*-value | *p*-value | *p* < .05 | *Cohen's d* |
|---|---|---|---|---|---|---|---|---|
| 1a | | | | | | | | |
| | Learning | HP vs. LP | -2.84 | 47 | 5.53 | < .001 | *** | 0.80 |
| | Test | Ego vs. LP | -2.14 | 47 | 3.03 | < .01 | ** | 0.44 |
| | Test | Allo vs. LP | -0.59 | 47 | 0.89 | .38 | n.s. | 0.13 |
| 1b | | | | | | | | |
| | Learning | HP vs. LP | -3.67 | 47 | 4.99 | < .001 | *** | 0.72 |
| | Test | Ego vs. LP | -0.57 | 47 | 1.19 | .24 | n.s. | 0.17 |
| | Test | Allo vs. LP | 0.72 | 47 | 1.47 | .15 | n.s. | 0.21 |
| 2a | | | | | | | | |
| | . | HP vs. LP | -0.42 | 47 | 0.86 | .39 | n.s. | 0.12 |
| | . | HP vs. prev HP | 0.0044 | 47 | 0.0073 | .99 | n.s. | 0.0011 |
| | . | Prev HP vs. LP | 0.42 | 47 | 0.58 | .56 | n.s. | 0.084 |
| 2b | | | | | | | | |
| | . | HP vs. LP | -0.53 | 47 | 1.28 | .21 | n.s. | 0.18 |
| | . | HP vs. prev HP | -0.20 | 47 | 0.34 | .74 | n.s. | 0.049 |
| | . | Prev HP vs. LP | 0.33 | 47 | 0.63 | .53 | n.s. | 0.091 |
| 2c | | | | | | | | |
| | . | HP vs. LP | -1.07 | 47 | 3.32 | < .01 | ** | 0.48 |

| | | | | | | | |
|---|---|---|---|---|---|---|---|
| . | HP vs. prev HP | -0.36 | 47 | 1.26 | 0.22 | n.s. | 0.18 |
| . | Prev HP vs. LP | 0.70 | 47 | 1.54 | 0.13 | n.s. | 0.22 |

[a] Difference in proportion of errors (%) between condition of interest and LP or prev HP condition.

**Supplementary table 4**

*Planned paired-sample t-test results for the RTs in Experiment 2c including target priming trials*

| Post-rotation | Planned comparison | Mean VSL effect (ms) | df | *t*-value | *p*-value | *p* < .05 | *Cohen's d* |
|---|---|---|---|---|---|---|---|
| First half | | | | | | | |
| | HP vs. LP | -24.66 | 47 | 7.35 | < .001 | *** | 1.061 |
| | HP vs. prev HP | -12.64 | 47 | 2.97 | < .01 | ** | 0.43 |
| | Prev HP vs. LP | -12.023 | 47 | 3.95 | < .001 | *** | 0.57 |
| Second half | | | | | | | |
| | HP vs. LP | -56.19 | 47 | 10.62 | < .001 | *** | 1.53 |
| | HP vs. prev HP | -56.43 | 47 | 8.02 | < .001 | *** | 1.16 |
| | Prev HP vs. LP | -0.77 | 47 | 0.22 | .83 | n.s. | 0.032 |

**Supplementary figure 1**

*Learned attentional suppression location across blocks in Experiment 1a*

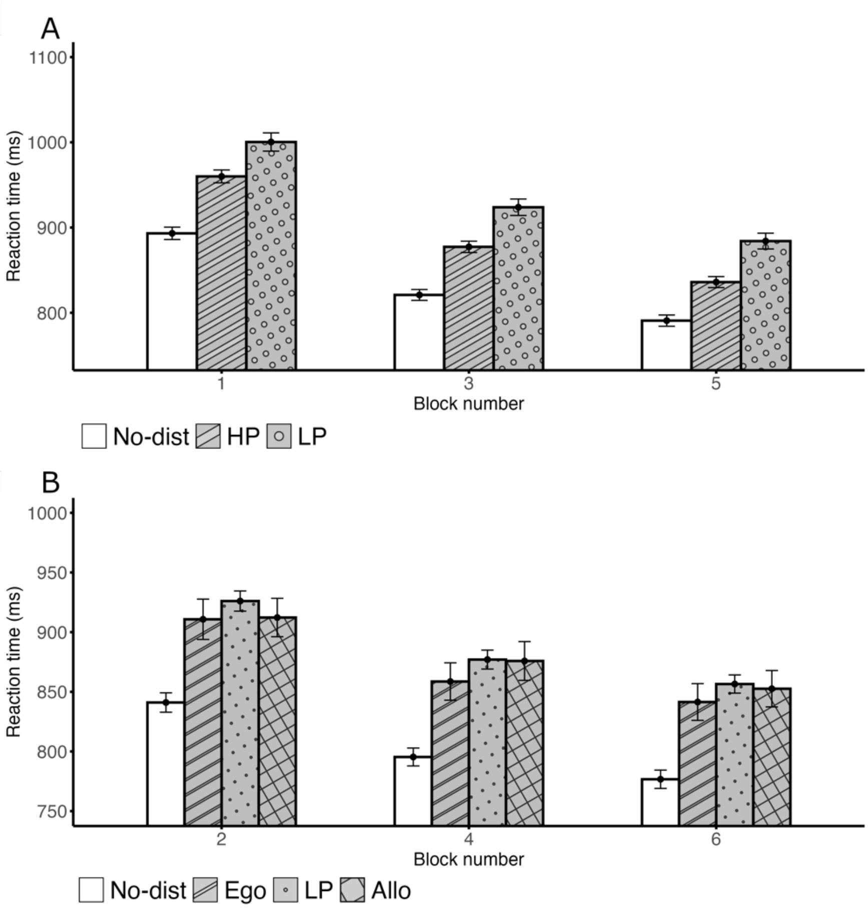


*Note.* Learning effects were evident in the first block of both the learning and test blocks. (A) Mean response times in the learning blocks. There was no significant interaction between Block number and Distractor condition (F (3.21, 150.75) = 2.11, *p* = .097, $n_p^2$ = .043, $BF_{excl}$ = 3.1) (B) Mean Response times in the test blocks. Similarly, no significant interaction was observed between Block number and Distractor condition (F(4.12, 193.7) = 0.31, *p* = .88 , $n_p^2$ = .007, $BF_{excl}$ = 180.6), indicating that suppression at the egocentric location remained consistent throughout the experiment.

**Supplementary figure 2**

*Learned attentional enhancement across blocks in Experiment 1b*

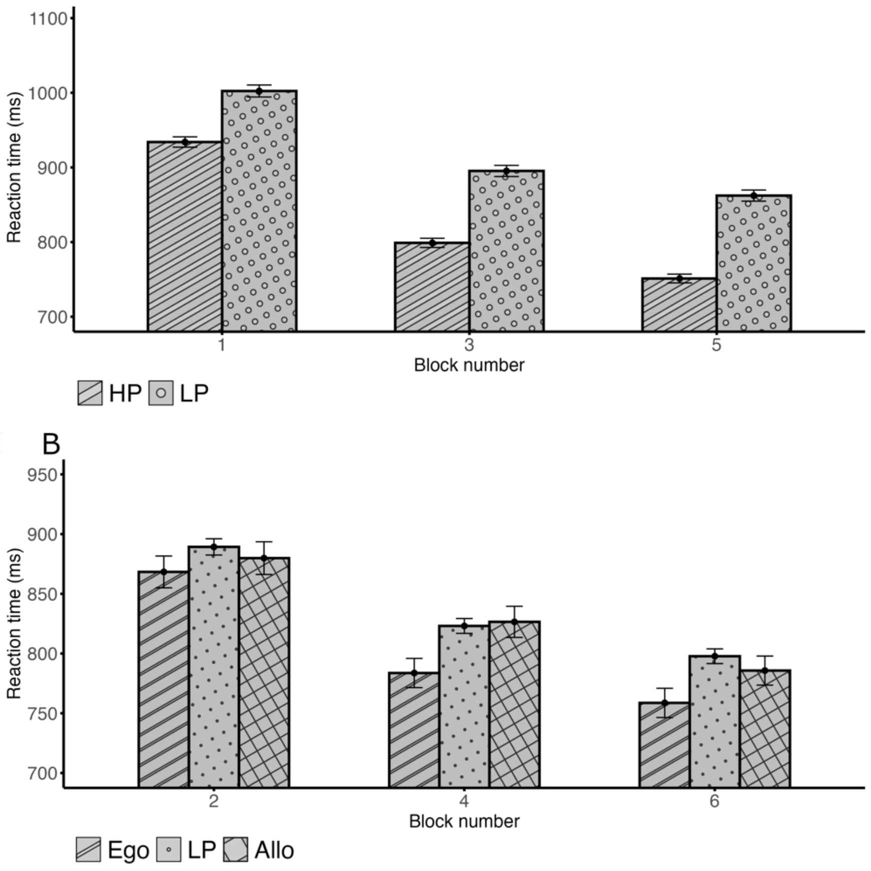


*Note.* Learning effects were evident in the first block of both the learning and test blocks. (A) Mean Response times in the learning blocks. In contrast to Experiment 1a, there was a significant interaction between Block number and Target condition (F (1.53, 71.97) = 14.01, *p* < .001, $n_p^2$ = .23). Despite this, each learning block demonstrated a significant difference between the HP and LP locations (all *t*'s > 8.0 and *p*'s < .001) (B) Mean Response times in the test blocks. Similar to the learning blocks, the test blocks showed a significant interaction between Block number and Target condition (F (3.31, 155.52) = 2.91, *p* = .032). Nevertheless, each block demonstrated a significant difference between the egocentric and LP locations (block 2: *t*(47) = 2.49, *p* = .016, *d* = 0.36; block 4: *t*(47) = 4.55, *p* < .001, *d* = 0.66; block 6: *t*(47) = 4.32, *p* < .001, *d* = 0.62), whereas the difference between the allocentric and

LP locations was not significant (block 2: $t(47) = 1.18$, $p = .25$, $d = 0.17$, $BF_{01} = 3.3$; block 4: $t(47) = 0.26$, $p = .79$, $d = 0.038$, $BF_{01} = 6.2$; block 6: $t(47) = 1.76$, $p = .085$, $d = 0.25$, $BF_{01} = 1.5$).

**Supplementary figure 3**

*Learned attentional suppression across blocks in Experiment 2a*

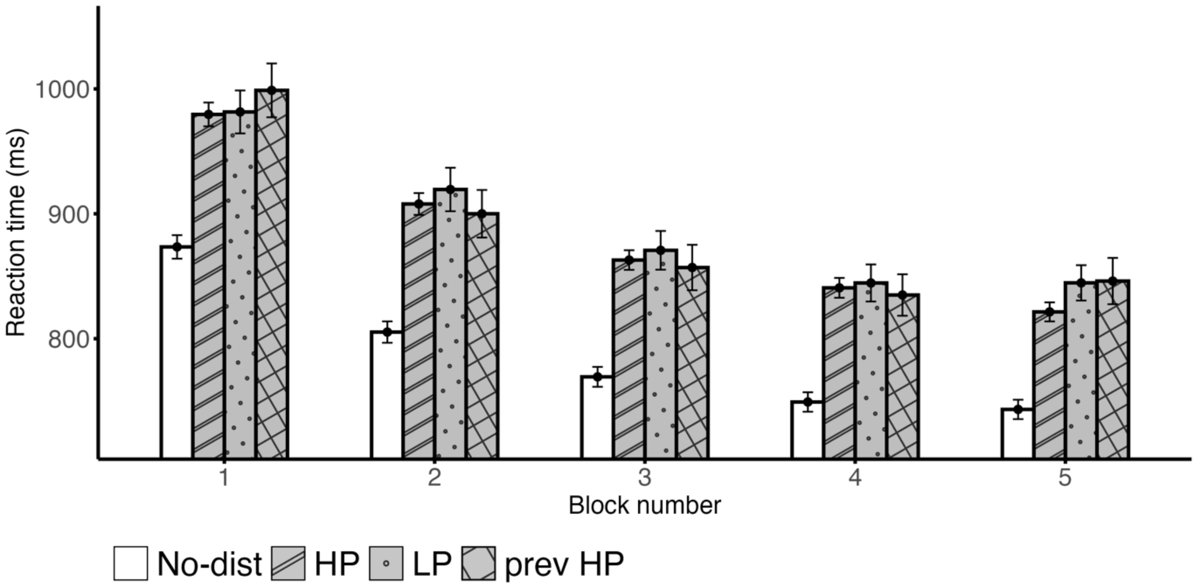


*Note.* Numerically the bias at the HP location appeared to be smallest in the first block and grew by the last block, but no significant interaction between Block number and Distractor condition was found (F (7.05, 331.57) = 1.55, $p = .15$, $n_p^2 = 0.032$, $BF_{excl} = 23.6$).

**Supplementary figure 4**

*Learned attentional enhancement across blocks in Experiment 2b*

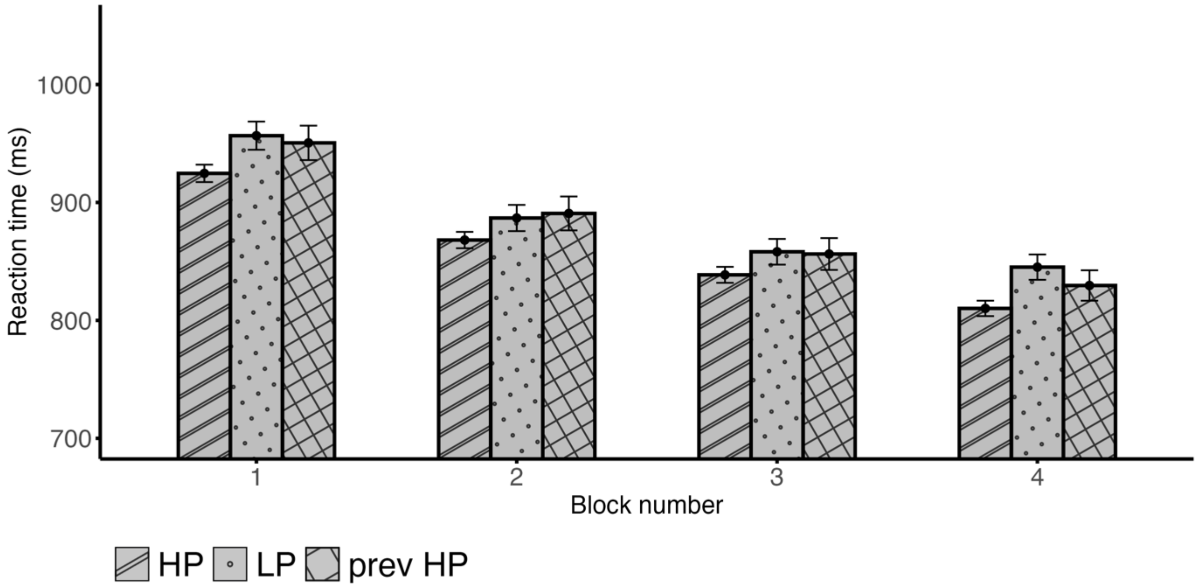


*Note.* The attentional bias toward the likely target location was evident from the first block, with no significant interaction between Block number and Target condition (F (6, 282) = 0.82, $p$ = .56, $n_p^2$ = 0.017, $BF_{excl}$ = 14.0).

**Supplementary figure 5**

*Learned attentional enhancement across blocks in Experiment 2c*

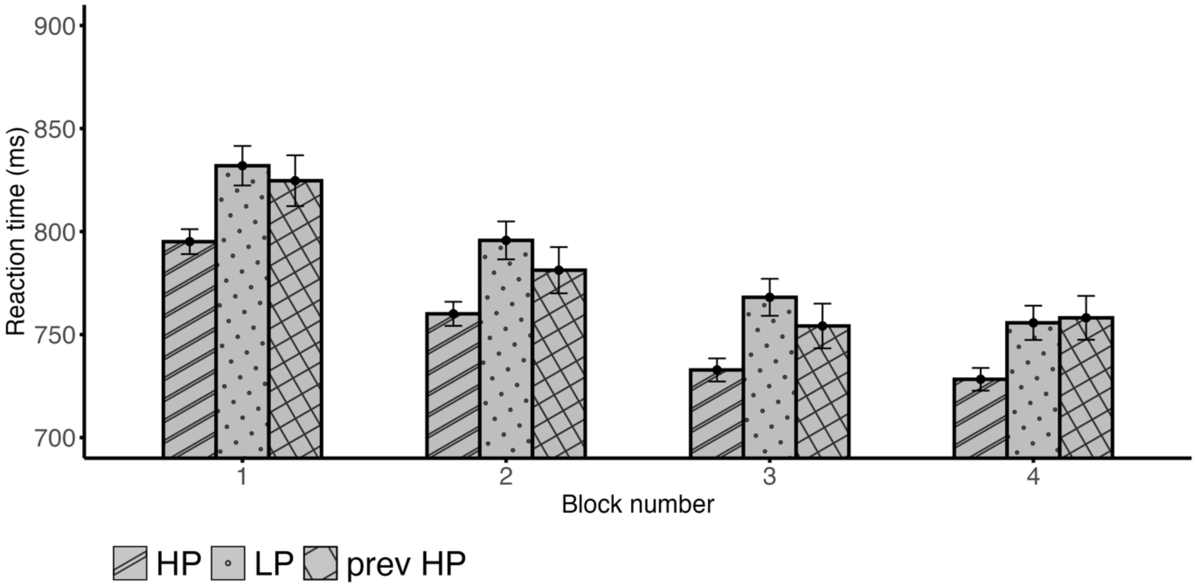


*Note.* The attentional bias toward the likely target location was evident from the first block, with no significant interaction between Block number and Target condition (F (4.6, 216.29) = 1.083, *p* = .37, $n_p^2$ = 0.023, $BF_{excl}$ = 7.6).
